# Inhibition of Cellular MEK/ERK Signaling Suppresses Murine Papillomavirus Type 1 Replicative Activities and Promotes Tumor Regression

**DOI:** 10.1101/2023.03.14.532042

**Authors:** Adrian J. Luna, Jesse M. Young, Rosa T. Sterk, Virginie Bondu, Fred A. Schultz, Donna F. Kusewitt, Huining Kang, Michelle A. Ozbun

## Abstract

Human papillomaviruses (HPVs) are a significant public health concern due to their widespread transmission, morbidity, and oncogenic potential. Despite efficacious vaccines, millions of unvaccinated individuals and those with existing infections will develop HPV-related diseases for the next two decades. The continuing burden of HPV-related diseases is exacerbated by the lack of effective therapies or cures for most infections, highlighting the need to identify and develop antivirals. The experimental murine papillomavirus type 1 (MmuPV1) model provides opportunities to study papillomavirus pathogenesis in cutaneous epithelium, the oral cavity, and the anogenital tract. However, to date the MmuPV1 infection model has not been used to demonstrate the effectiveness of potential antivirals. We previously reported that inhibitors of cellular MEK/ERK signaling suppress oncogenic HPV early gene expression *in vitro*. Herein, we adapted the MmuPV1 infection model to determine whether MEK inhibitors have anti-papillomavirus properties *in vivo*. We demonstrate that oral delivery of a MEK1/2 inhibitor promotes papilloma regression in immunodeficient mice that otherwise would have developed persistent infections. Quantitative histological analyses revealed that inhibition of MEK/ERK signaling reduces E6/E7 mRNAs, MmuPV1 DNA, and L1 protein expression within MmuPV1-induced lesions. These data suggest that MEK1/2 signaling is essential for both early and late MmuPV1 replication events supporting our previous findings with oncogenic HPVs. We also provide evidence that MEK inhibitors protect mice from developing secondary tumors. Thus, our data suggest that MEK inhibitors have potent anti-viral and anti-tumor properties in a preclinical mouse model and merit further investigation as papillomavirus antiviral therapies.

**Significance Statement:** Persistent human papillomavirus (HPV) infections cause significant morbidity and oncogenic HPV infections can progress to anogenital and oropharyngeal cancers. Despite the availability of effective prophylactic HPV vaccines, millions of unvaccinated individuals, and those currently infected will develop HPV-related diseases over the next two decades and beyond. Thus, it remains critical to identify effective antivirals against papillomaviruses. Using a mouse papillomavirus model of HPV infection, this study reveals that cellular MEK1/2 signaling supports viral tumorigenesis. The MEK1/2 inhibitor, trametinib, demonstrates potent antiviral activities and promotes tumor regression. This work provides insight into the conserved regulation of papillomavirus gene expression by MEK1/2 signaling and reveals this cellular pathway as a promising therapeutic target for the treatment of papillomavirus diseases.

## Introduction

Human papillomavirus (HPV) infections are highly prevalent and cause a variety of epithelial neoplasms (i.e., tumors) ranging in severity from benign cutaneous warts to precancerous intraepithelial neoplasia to cancers (1). Over 200 HPV genotypes have been identified and sequenced (2) with low-risk mucosotropic HPVs causing 90% of anogenital warts and all laryngeal papillomas and high-risk HPV infections etiologically linked to anogenital and oropharyngeal cancers (3). Despite effective prophylactic vaccines, millions of unvaccinated people and those with existing infections are projected to develop HPV-related diseases for the next two decades (4) and beyond (5). Current HPV treatments involve lesion excision or ablation, cryotherapy, chemotherapy, and radiation, each risking long-term morbidities with reduced quality of life (6–9). Additionally, recurrence rates for some treatments can be >50% (10, 11). To date, there are no effective antiviral HPV therapies and no cure for most HPV infections.

Like all PVs, HPVs are highly host specific and cannot be productively transmitted to animals. Although this has inhibited detailed studies of HPV pathogenesis and immunology, experimental infections using a variety of vertebrate animal PVs have provided valuable models for understanding viral gene functions, tissue tropism, regression and persistence, cancer progression, vaccine efficacy, and potential therapeutic agents (12). Despite differences in epithelial tropisms and disease manifestations, mammalian PVs have similar genome organization, protein functions, and reliance on epithelial differentiation for productive replication (1). The murine PV type 1 (MmuPV1) mouse model is a major advancement in preclinical models of PV-induced infection and disease (13). Whereas mouse models are frequently used to identify and test candidate interventions and the MmuPV1 should be a valuable preclinical model, to our knowledge, this model has not been used to demonstrate the efficacy of antiviral agents.

In all mammalian PVs, tumorigenesis is primarily driven by the E6 and E7 proteins (14, 15). Although E6 and E7 from different PV genera interact with different cellular targets, E6 and E7 expression reprograms cells in the para- and supra-basal layers to maintain cell cycle progression and proliferative capacity (14, 16, 17). This promotes a cellular environment where viral genomes are amplified for packaging by the capsid proteins into new virions (16). Additionally, E6 and E7 proteins play critical roles in viral genome maintenance and amplification to support productive PV infections (18, 19). As E6 and E7 proteins are the key factors for PV persistence and oncogenesis, they are frequently considered to be ideal therapeutic targets (20, 21). Genetic knockdown of oncogenic HPV E6 and E7 expression restores cell cycle checkpoint (e.g., p53 and pRb) functions, inhibits cell proliferation, and leads to cellular apoptosis and/or senescence (22–30). Yet, genetic approaches have not been robustly adaptable to human clinical studies. A handful of small molecules have been investigated for their abilities to inhibit PV protein activities (31), but none so far show promise *in vivo*.

We recently reported that cellular MEK/ERK signaling is a key modulator of high-risk HPV E6/E7 transcription (32). The RAS-RAF-MEK-ERK signaling pathway is evolutionarily conserved, has been extensively investigated in mammalian cells, and is the best understood signaling cascade in tumor cell biology (33). MEK signaling is tightly controlled in normal epithelium where proliferative signals from growth factor receptors are restricted to promote epithelial homeostasis, including cellular contact inhibition and differentiation (34–37). Thus, MEK signaling is normally limited to the lower epithelial cell layers. MEK signaling is effected by the MEK1 and MEK2 isoforms, which are structurally similar and share activation mechanisms. Upon activation by RAF, they function as dual-specificity tyrosine and serine/threonine kinases, phosphorylating their only known *in vivo* substrates ERK1 and ERK2 on specific tyrosine and threonine residues (38). MEK1 and MEK2 exhibit extensive functional redundancy and form functional homo- or hetero-dimers *in vivo* (39). However, increasing evidence suggests that MEK1 and MEK2 have distinct activities that may be controlled by specific modulation of their activity levels (40, 41), differential binding to cellular proteins (42–46) and disparate outcomes when the genes are knocked out in mice (39, 47, 48). Despite an incomplete understanding of the distinct functions of MEK1 and MEK2, particularly in the epithelium, MEK/ERK activation has long been recognized to play a key role in cell proliferative diseases and neoplastic progression. For example, p-ERK1/2 levels are increased in the upper epithelial layers in several proliferative skin diseases, including HPV-driven cervical neoplasia (49). We found, in a cohort of 580 biopsies from normal cervix, low-, and high-grade cervical intraepithelial neoplasia (CIN), a significant correlation (p<0.001, Ξ^2^=212.7) between increasing levels of p-ERK and E7 protein activity during disease progression (32). Mechanistically, using high-risk HPV-infected 3D organotypic (raft) epithelial tissues and monolayer cultures, we showed that MEK-induced phosphorylation of ERK1/2 robustly governs HPV E6/E7 mRNA transcription. Importantly, we demonstrated that small molecule inhibitors of MEK and ERK potently inhibit HPV E6/E7 transcription, at least in part by blocking the activation of activator protein-1 (AP-1) transcription factors (32) whose binding sites are conserved in the long control region (LCR) enhancer/promoter of all PV genomes (50).

The combined conservation of PV genome structure, epithelial differentiation-dependent PV gene regulation, and MEK/ERK signaling in mammalian epithelium prompted us to hypothesize that inhibitors of MEK signaling would suppress E6/E7 gene transcription and tumor growth in the preclinical MmuPV1-induced tumor model that recapitulates key aspects of HPV-induced tumorigenesis (13). We investigated two FDA- and European Medicines Agency-approved MEK inhibitors, trametinib and cobimetinib (51). Both drugs have similar *in vitro* inhibitory concentration 50 (IC_50_) efficacies for preventing activation of MEK1 (0.7 nM and 0.95 nM, respectively) and trametinib is equivalently inhibitory of the MEK2 isoform (IC_50_ = 0.9 nM). However, cobimetinib is ≈200-fold less potent against MEK2 (IC_50_ = 199 nM) (51). After inducing tumors on the tails of Hsd:Athymic-*FoxN1^nu^* mice that lack T cells, mice were treated orally with drug dosages based on previous studies (51). Both drugs effectively inhibited tumor growth, but trametinib led to significant tumor regression beginning about 1 week post treatment initiation. Furthermore, the MEK inhibitors prevented overt lesion formation at secondary sites, suggesting diminished MmuPV1 transmission. Both drugs display mechanisms of action consistent with reduced viral gene expression and genome replication and suppressed cellular proliferation as expected from our prior *in vitro* study (32). Our results strongly suggest that both MEK1 and MEK2 are important for PV tumorigenesis, and that further investigations of MEK1/2 inhibitors are warranted for the treatment of PV-induced diseases in humans and animals.

## Results

### MEK Inhibitors Suppress MmuPV1-Induced Tumor Growth and Reduce Papillomatous Morphology

After MmuPV1 inoculation and 6 weeks of papilloma growth, mice were randomized into treatment groups; the average tumor volume of each group was ≈40 mm^3^ (Fig. 1A) with no significant size differences among the groups (*SI Appendix* Table 1). Mice were orally administered either vehicle, the MEK1 inhibitor cobimetinib (7.5 mg/kg or 15 mg/kg), or the MEK1/2 inhibitor trametinib (1.0 mg/kg or 2.0 mg/kg) every 2 days. As visualized in Fig. 1B, C and detailed in *SI Appendix* Table 1, tumor volumes significantly differed among the five groups after 8 days of treatment (p=0.042) and the difference became increasingly pronounced as time passed. Over the 30-day treatment period, tumors of vehicle-treated mice increased in size by an average of 2.59-fold, resulting in a final average tumor volume of 101.7 mm^3^. Comparatively, mice treated with 7.5 mg/kg or 15 mg/kg of cobimetinib demonstrated significantly impaired papilloma growth resulting in final average tumor volumes of 42.4 mm^3^ (58% TGI) and 45.4 mm^3^ (55% TGI), respectively (*SI Appendix* Table 1). More strikingly, trametinib treatment resulted in papilloma regression over the same period, reducing tumor volumes to an average of 25.0 mm^3^ for 1 mg/kg trametinib and 25.2 mm^3^ for 2 mg/kg trametinib (both 75% TGI; *SI Appendix* Table 1). Pairwise comparisons revealed significant differences between vehicle-treated and each of the drug treatment groups (adjusted p-value <0.05), but no significant differences when comparing each of the four groups that received the MEK inhibitors (*SI Appendix* Table 2). Similarly, the rates of tumor volume change (mm^3^/day) among the treatment groups estimated by linear mixed-effects model analysis showed that the vehicle-treatment group significantly increased over time while the trametinib-treated groups significantly decreased, and the cobimetinib-treated groups did not have a significant change (*SI Appendix* Table 3). Furthermore, trametinib demonstrated significant superiority over cobimetinib in promoting tumor regression as demonstrated by comparing the rates of tumor volume change among the MEK inhibitor treated groups (p-value = 0.0334; *SI Appendix* Table 4). The visual characteristics of the tail papillomas on mice treated with MEK inhibitors were distinct compared to papillomas on vehicle-treated mice (Fig. 1D; *SI Appendix* Fig. S1). Specifically, vehicle-treated mice bore florid and irregular papillomas. Whereas mice treated with both cobimetinib doses exhibited smaller tumors with less florid appearance, mice treated with both trametinib doses generally displayed more flattened and smoother papillomas (Fig. 1D; *SI Appendix* Fig. S1). Note that only one animal, in the high dose trametinib treatment group, showed significant weight loss suggesting toxicity (*SI Appendix* Fig. S1F-H). These data indicate that MEK inhibitors have profound TGI effects *in vivo* using the MmuPV1 preclinical model; compared to the MEK1 inhibitor cobimetinib, the MEK1/2 inhibitor trametinib, demonstrated superior TGI.

**Figure 1.**
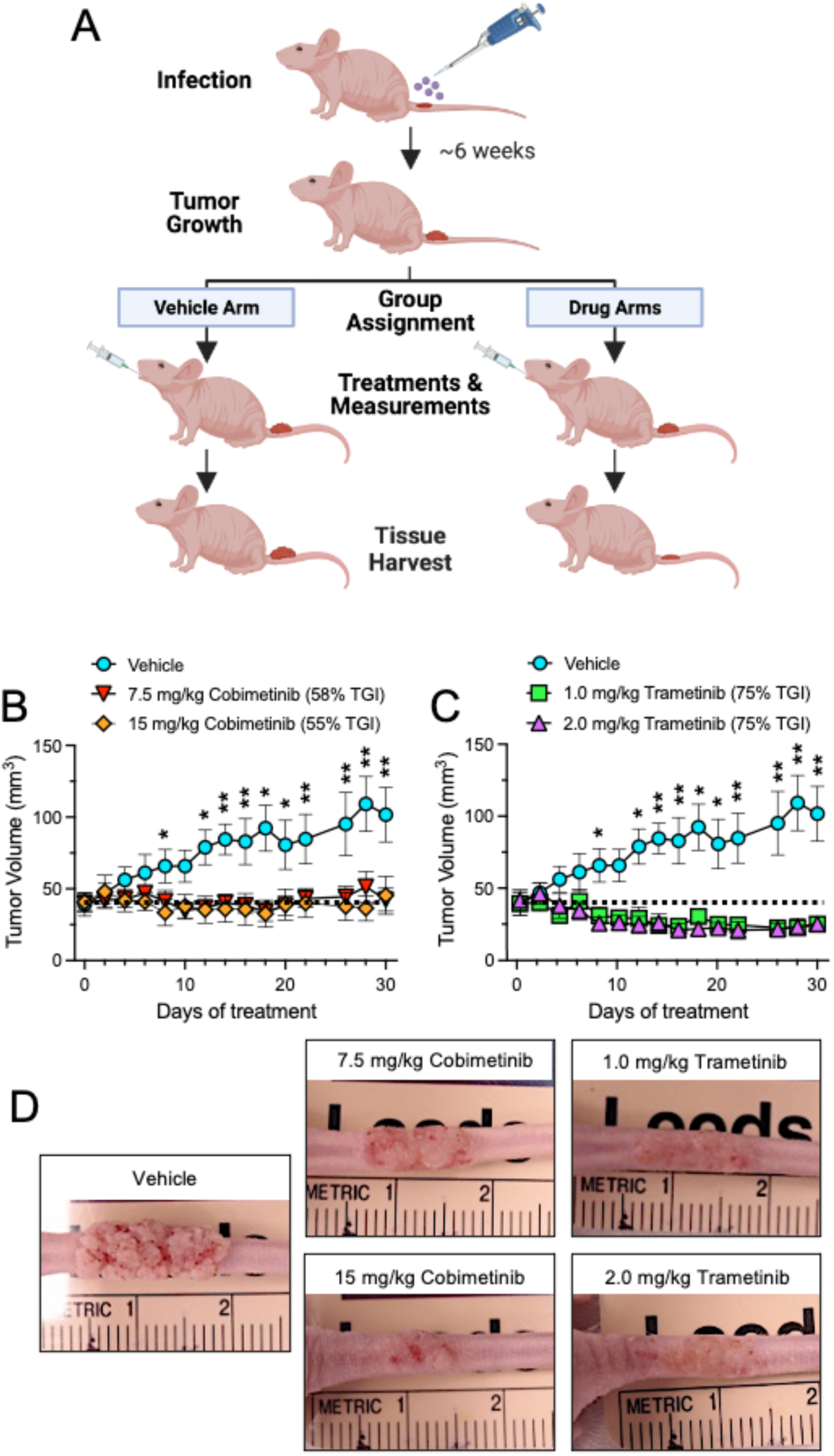
MEK inhibition impairs MmuPV1-induced papilloma growth. (A) Immediately after epithelial tail scarification, female Hsd:Athymic Nude-*Foxn1^nu^* mice were exposed to 2×10^8^ vge of MmuPV1. Papillomas were allowed to grow for 6 weeks before assigning the animals into vehicle or treatment groups with similar average tumor volumes (n=6 animals per group). Mice were treated by oral gavage treatment every 48h before harvesting the tumor-bearing tail sections on day 30 post treatment. Image created with BioRender.com. (B, C) Mean tumor volumes tracked over the treatment period with (B) cobimetinib or (C) trametinib (the vehicle group was the same for both drugs). The dotted horizontal line marks the mean starting volumes. Data were analyzed comparing all treatment groups together using ANOVA with the generalized least square method (*p<0.05 and **p<0.01; *SI Table 1*); the treatments were graphed separately for clarity. Error bars represent SE. Mean TGI for the MEK inhibitors were 58% and 55% for the 7.5 mg/kg and 15 mg/kg cobimetinib, respectively and 75% for both doses of trametinib. (D) Representative images of the MmuPV1 induced papillomas for each treatment group at day 30 post treatment initiation.

### MEK Inhibitor Effects on Tissue Morphology and Cellular Biomarkers

Serial, sagittally sectioned tails were analyzed histologically after H&E staining, focusing on the mice treated with vehicle and those treated with the higher doses of the two MEK inhibitors (Fig. 2A; *SI Appendix* Fig. S2). Papillomas from vehicle-treated mice showed extensive fibrillary projections throughout the tumor cross-sections. In drug-treated mice, particularly those treated with the MEK1/2 inhibitor, trametinib, the extent of papillary projections was considerably reduced (Fig. 2A). Consistent with this observation, morphometric analyses of all tumors revealed significantly reduced epithelial thickness in the papillomas from mice treated with the MEK inhibitors compared to vehicle-treated mice (Fig. 2B). Tumor sections from all three groups revealed dysplastic epithelium and hyperkeratosis, albeit with noticeable differences depending upon the treatment (*SI Appendix* Fig. S2). Koilocytes, epithelial cells characterized by perinuclear haloes (cytoplasmic vacuolation) surrounding condensed nuclei, are pathognomonic of HPV infections (52) and were scattered irregularly throughout the differentiating epithelium in all MmuPV1 lesions.

**Figure 2.**
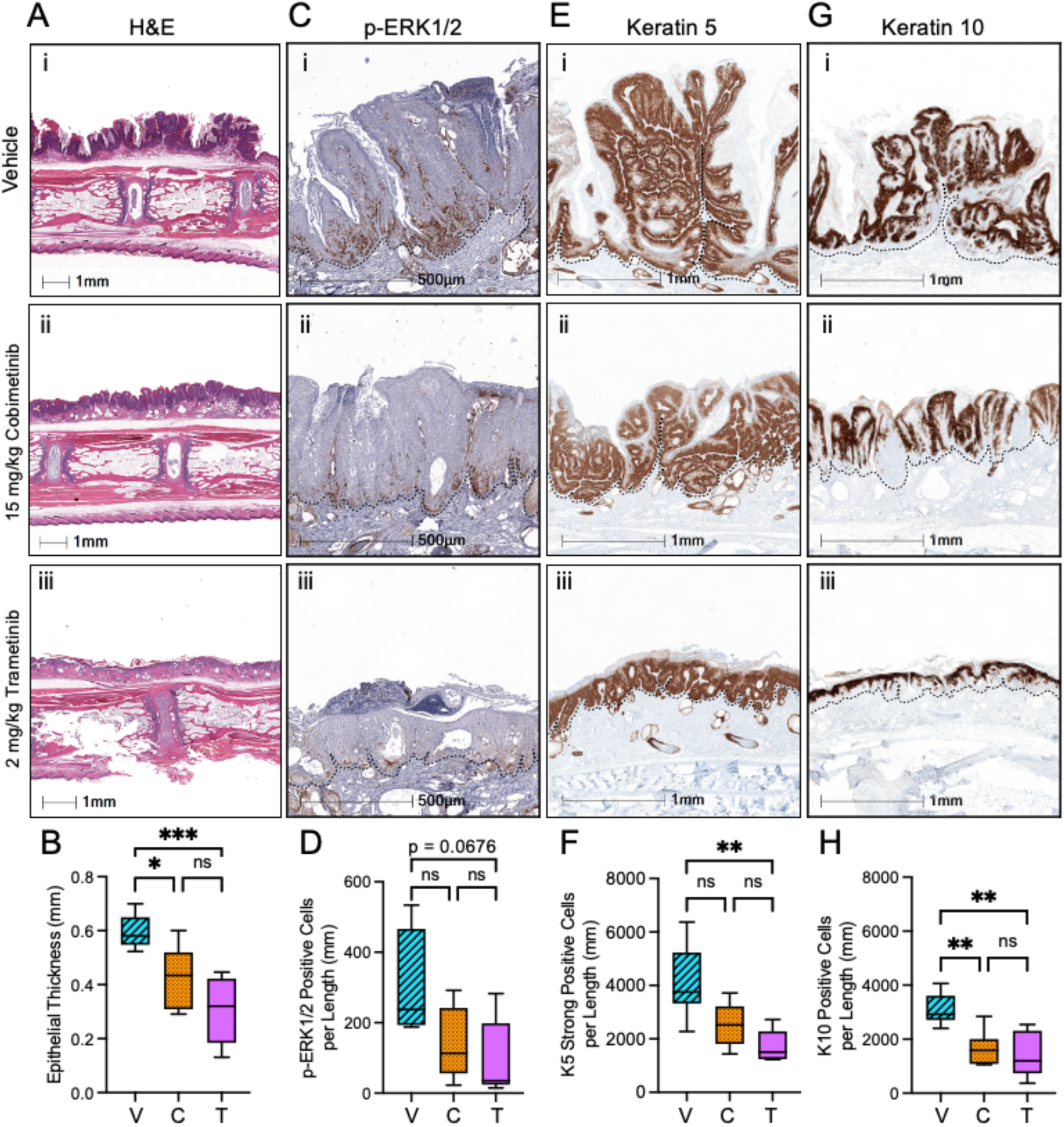
Histological and immunohistochemical analysis of MmuPV1-induced papillomas. Representative images of papilloma histology from mice treated with vehicle (i),15 mg/kg cobimetinib (ii), or 2.0 mg/kg trametinib (iii). Tails were fixed with paraformaldehyde, paraffin embedded, sectioned, and stained: (A) H&E; (C) IHC for p-ERK1/2; (E) IHC for keratin 5 (K5); (G) IHC for keratin 10 (K10); dotted lines mark the basement membrane. (B, D, F, H) Morphometric quantification of tail tumors from each mouse (4-5 per group) for papilloma thickness (height, B); p-ERK1/2 (D), K5 (F), and K10 (H) IHC staining expressed as positive cells per tumor length (mm); vehicle (V), 15 mg/ml cobimetinib (C), 2.0 mg/kg trametinib (T). Data were analyzed using one-way ANOVA with Tukey’s multiple comparison test (*p<0.05, **p<0.01; ***p<0.001).

The MEK inhibitors suppressed the levels of phosphorylated-ERK1/2 (p-ERK1/2), the only recognized signaling effector of MEK, in the tumors (Fig. 2C-D); moreover, p-ERK1/2 levels have a strong correlation with tumoral epithelial lesion thickness (r=0.7151, p=0.00185; *SI Appendix* Fig. S5A). p-ERK1/2 was most prominent in the papillomas of vehicle-treated mice, localizing primarily to basal and lower suprabasal layers of all papillomas. These data indicate that the systemically delivered MEK inhibitors reached the tail epithelium to suppress MEK/ERK signaling. Further, the findings suggest that MEK signaling is an important contributor to papilloma growth and maintenance in the MmuPV1 mouse model.

As MEK/ERK signaling is integral to epithelial proliferation and differentiation (53), we evaluated the papillomas for differentiation-associated epithelial markers, keratin 5 (K5) and keratin 10 (K10). K5 pairs with K14 and these proteins are characteristic of mitotically active, undifferentiated basal cells of the epithelium; their expression is downregulated as cells begin to differentiate (54). In contrast, K10 is an established early marker of keratinocytes entering terminal differentiation (55). As expected, strong K5 staining was prominent in the cells nearest the basement membranes of all tumors. The vehicle-treated mice displayed an expansive network of undifferentiated cells throughout the thickness of the fibrillary papillomas (Fig. 2E; *SI Appendix* Fig. S3A-D), as previously reported for MmuPV1 tail lesions (17). However, MEK inhibitors counteracted basal cell expansion and the number of K5-positive cells was reduced (Fig. 2F) and were correlated with p-ERK1/2 levels (r=0.5947, p=0.0249) and strongly correlated with lesion thickness (r=0.7641, p=0.000569; *SI Appendix* Fig. S5B-C, respectively). Likewise, the number of differentiating K10-positive cells was significantly lower in the smaller papillomatous lesions of the drug-treated mice compared to those in vehicle-treated mice (Fig. 2G-H). Interestingly, there were no significant differences amongst the treatment groups in the percentages of lesional cells that expressed p-ERK1/2, K5, or K10 (*SI Appendix* Fig. S3E-G). These findings indicate that while MEK inhibition and suppressed p-ERK1/2 signaling reduced the number of proliferative cells, there were no net changes in the overall proportion of cells proliferating or differentiating within the papillomas.

Infection with MmuPV1 or oncogenic HPVs leads to increased expression of Ki67, a basal cell proliferation marker (56–59) and Sundberg et al. showed similar staining patterns between Ki67 and K5 in MmuPV1-induced tail tumors (56). Furthermore, MEK inhibition is known to reduce Ki67 staining in epithelial tumors (60, 61). Although we attempted to detect Ki67 by immunohistochemistry (IHC) in the tumor sections, we were unable to obtain quantifiable Ki67 staining. This is likely because tissue decalcification has been found to impair the detection of nuclear biomarkers (62).

### MEK Inhibitor-Induced Papilloma Regression does not Involve T Cells or Neutrophils

Whereas various data indicate that CD4+ and CD8+ T cells play a major role in papillomavirus-induced lesion regression (56, 63–66), there also are indications that neutrophils might contribute to MmuPV1 disease control in immunocompetent models (67). As the Hsd:Athymic-*FoxN1^nu^* mice used in this investigation lack CD4+ and CD8+ T cells but have an intact innate immune system, we investigated if MEK inhibition resulted in increased neutrophil infiltration in the MmuPV1-induced papillomas. Quantification of tumor-proximal myeloperoxidase staining revealed no significant differences in neutrophil infiltration amongst treated and control papillomas (Fig. 3A-B). Thus, the data suggest that MEK inhibitors suppressed tumor growth and trametinib promoted regression *via* a mechanism independent of T cells and neutrophils.

**Figure 3.**
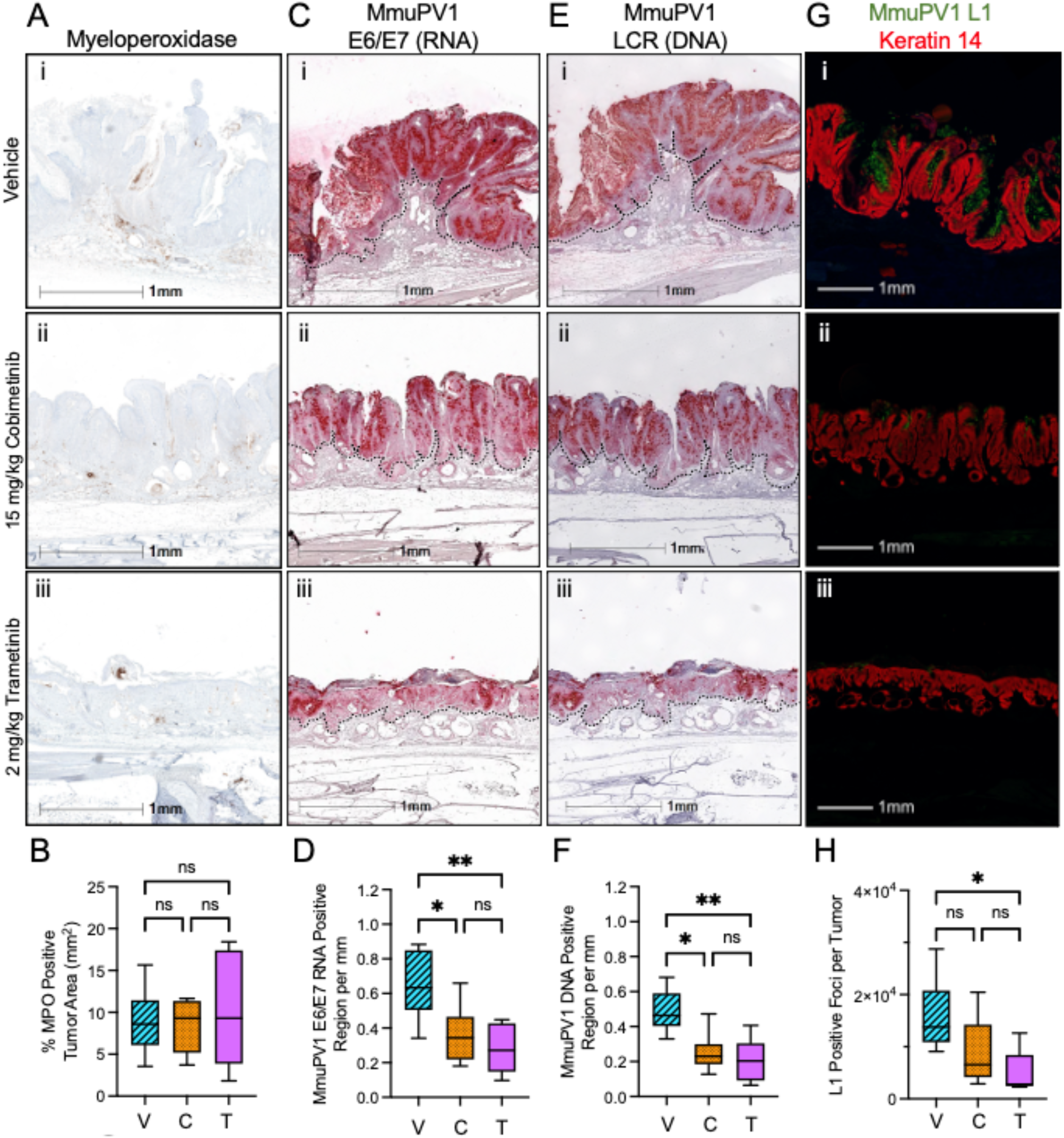
Histological analysis of immune and viral markers in MmuPV1-induced papillomas. Representative images of fixed histological papilloma sections for mice treated with vehicle (i),15 mg/kg cobimetinib (ii), or 2.0 mg/kg trametinib (iii). Tissues were stained *via*: (A) IHC for neutrophil marker, myeloperoxidase (MPO); (C) ISH for MmuPV1 E6/E7 mRNA; (E) ISH for MmuPV1 genomic DNA (note serial sections in panels C and E for comparison); (G) IF for MmuPV1 L1 protein (green) and keratin 14 (red). The brightness of the photos in panel G was enhanced equivalently to better illustrate L1 staining. (B, D, F, H) Morphometric quantification of each tumor section (vehicle, V; 15 mg/ml cobimetinib, C; 2.0 mg/kg trametinib, T) on raw images (n = 5-6/group) for the percent of MPO positive cells in the tumor (B), region of positive E6/E7 ISH staining per mm of papilloma (D), region of positive LCR ISH staining per mm of papilloma (F), and number of L1 positive foci per tumor (H). Data were analyzed using one-way ANOVA with Tukey’s multiple comparison test (*p<0.05, **p<0.01).

### The MmuPV1 Vegetative Replication Cycle is Dependent on MEK Signaling

Our recent publication demonstrated that HPV early gene expression relies on MEK/ERK signaling and that MEK1/2 inhibition results in reduced viral early gene transcription and protein expression (32). As robust PV genome amplification leading to late gene expression depends on sufficient PV early gene expression (18, 19, 68), we determined the effect of MEK inhibition on key aspects of the complete MmuPV1 replicative cycle. We first strove to quantify the tumor levels of E6/E7 oncogene mRNA, which should be present predominantly in the cytoplasm of infected epithelium (Fig. 3C). Despite various DNase treatment methods to remove viral genomes, the E6/E7 probes consistently bound to nuclear targets, particularly evident in the vehicle-only tumors (*SI Appendix* Fig. S4A). Thus, the data in Fig. 3C-D likely represent detection of MmuPV1 genomes and E6/E7 mRNAs; yet the morphometric analyses demonstrate the MEK inhibitors significantly reduced the levels of MmuPV1 nucleic acids in the tumors. Consistent with this interpretation, the *in situ* hybridization (ISH) probe specific to the MmuPV1 noncoding long control region (LCR), revealed ISH staining patterns with obvious overlap in serial sections (Fig. 3, compare panels C and E; *SI Appendix* Fig. S4). Morphometric quantification of ISH data showed overall lower levels of LCR-targeted viral genomes compared to the E6/E7 ISH results in each treatment group (Fig. 3F and D, respectively). There was a strong correlation between E6/E7 ISH and LCR ISH (r=0.9263, p<0.00001; *SI Appendix* Fig. S5J). Furthermore, both RNA ISH and DNA ISH levels had strong correlations with p-ERK1/2 levels (r=0.8303, r=0.7421, both p<0.001; *SI Appendix* Fig. S5E-F, respectively). Together, the data indicate significantly decreased viral genome levels and imply reduced E6/E7 mRNAs in tumors from mice receiving the MEK inhibitors. Lastly, IF detection of MmuPV1 major capsid protein L1 along with K14 staining of undifferentiated basal cells in the tumor sections revealed significantly fewer L1-positive foci in the differentiating layers of tumors of the cobimetinib and trametinib treatment groups (Fig. 3G,H). Additionally, E6/E7 and viral LCR ISH were strongly correlated with L1-foci (r=0.7245, p<0.0015 and r=0.8411, p<0.0001; *SI Appendix* Fig. S5L-M, respectively). These data clearly support the idea that the MEK inhibitors suppressed the MmuPV1 replicative cycle *in vivo*.

### MEK Inhibitor Treatment Reduces MmuPV1 Transmission *in vivo*

Over the treatment course, we observed that all (6/6) vehicle-treated mice developed papillomas on their muzzles (Fig. 4A; Table 1). In contrast, the only mouse from the MEK inhibitor treatment cohorts to develop a visible facial wart (1/24) was in the group receiving the lower dose (7.5 mg/kg) of cobimetinib (Table 1). This mouse was co-housed with other animals receiving cobimetinib (i.e., it was not co-housed with vehicle-treated mice). H&E staining and E6/E7 ISH confirmed that the facial lesions arose from MmuPV1 infections (Figure 4B). As the snouts were not experimentally scarified or infected, the secondary muzzle-arising infections were likely transmitted from tail grooming. The most obvious explanation for lack of secondary muzzle infections in MEK inhibitor-treated mice is that their tail papillomas produced fewer virions due to suppressed viral replication as shown in Fig. 3C-H. We reported that MEK/ERK signaling stimulates HPV early gene expression *via* activation of cellular fos/jun family transcription factors (32), which bind to AP-1 binding sites that are highly conserved in papillomavirus LCRs (69). Thus, it is also possible that MEK inhibition would result in lower viral early gene transcription after infection-mediated delivery of viral genomes to the cell nucleus. To test this prospect, we exposed HaCaT keratinocytes to MmuPV1 virions in the presence of increasing doses of trametinib. Indeed, MEK inhibition resulted in significantly decreased levels of spliced MmuPV1 E1^E4 mRNAs compared to vehicle treated cells (Fig. 4C). Combined, our data suggest that MEK inhibition is likely to impede new MmuPV1 infections by both reducing virion shedding from existing lesions and by suppressing early viral transcription during infection establishment at new sites.

**Figure 4.**
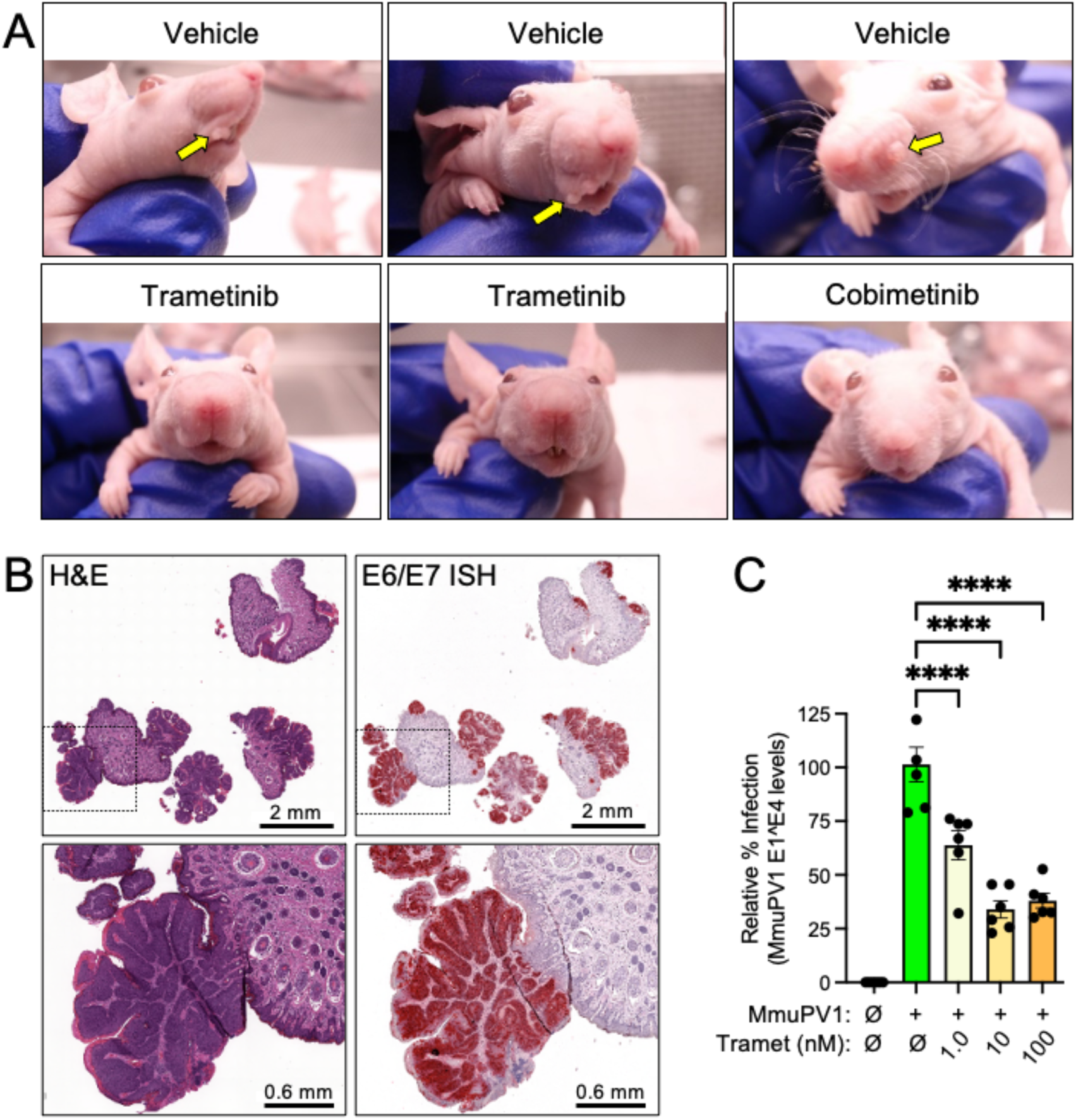
MEK inhibition provides protection against MmuPV1 transmission to secondary epithelial sites. (A) Mice that were co-housed throughout the 30-day treatment period; arrows indicate papillomas that developed on the muzzles of vehicle-treated mice. (B) Histological images of facial papillomas harvested on day 30 post treatment. Tissue sections were stained with H&E or by MmuPV1 E6/E7 ISH. Smaller boxes in upper panels indicate regions of increased magnification (lower panels). (C) Relative MmuPV1 infection in untreated cells compared to those treated with increasing concentrations of trametinib. Data summarize three independent infections analyzed by one-way ANOVA with Dunnett’s multiple comparisons test (**** p<0.0001).

**Table 1.**
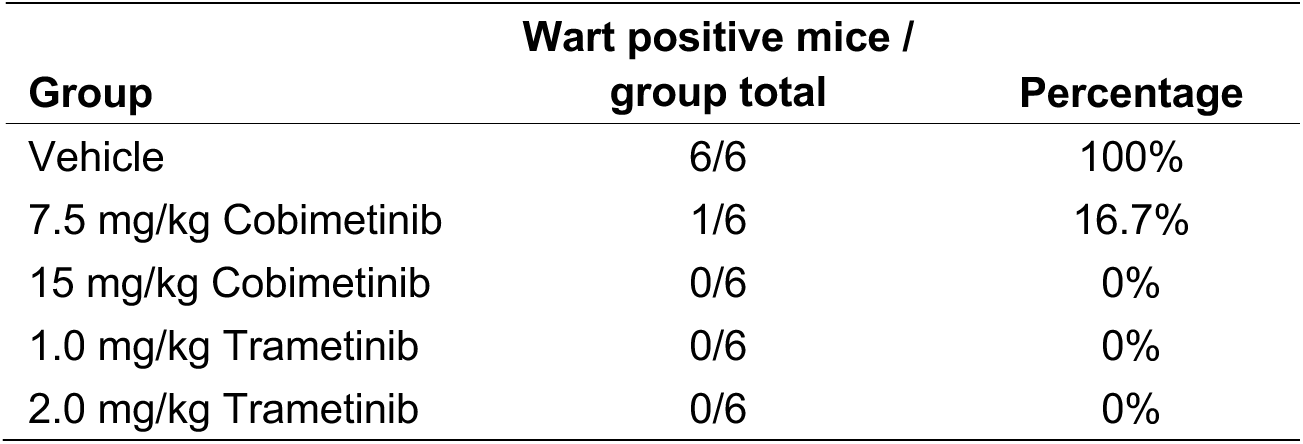
Summary of animals developing visible muzzle warts

## Discussion

Herein we demonstrate both the important role for MEK1/2 signaling in PV-induced tumorigenesis and the potent antiviral and tumor suppressive properties of MEK1/2 inhibitors in a preclinical, natural infection model of HPV disease. This work confirms and extends our recent study showing that inhibiting MEK/ERK signaling has antiviral effects, specifically reducing early gene transcription and oncoprotein functions, against oncogenic HPVs in three dimensional organotypic epithelial tissue cultures (32). The conservation of MEK/ERK signaling in mammalian epithelium coupled with the conservation of MEK/ERK activated AP-1 transcription factor binding sites in the LCRs of animal PVs indicates that this pathway is both critical for controlling PV early gene transcription and a prime target for PV antivirals. Thus, our work suggests that targeting this pathway is likely to have antiviral effects for all mammalian PV infections. Importantly, in our prior work and in this study, the antiviral and tumor suppressive effects of MEK inhibitors on PV infections occurred in the absence of T cells, which has significant clinical implications for treating PV infections in individuals lacking functional T cells.

An important aspect of our study that is particularly relevant to treating pathogenic HPV induced lesions is the fact that we initiated treatments only after MmuPV1 infection induced clinically evident papillomas (average of ≈40 mm^3^ in each cohort) six weeks post inoculation. Most preclinical studies designed to test candidate therapies against PV disease begin treatments prior to pathologically confirmed disease. For example, a study investigating the antiviral effects of dihydroartemisinin (DHA, an active metabolite of artesunate) showed topical treatment could inhibit the development of mucosal tumors in two of three animals exposed to canine oral papillomavirus (70). However, the treatment was initiated 24h after viral challenge and ≈3 weeks prior to tumor development in the vehicle-treated animals. More recently, DHA was tested following MmuPV1 challenge in the murine anal or cervicovaginal tract; however, neither the persistence of viral infections nor the presence of disease was verified at these sites prior to treatment initiation (71). Even so, DHA failed to reduce disease development at either site. This was surprising given that topical vaginal delivery of artesunate led to histologic regression in 19/28 (67.9%) of immunocompetent women with HPV-induced CIN grade 2/3 and HPV clearance in 9/19 (47.4%) subjects whose lesions underwent histologic regression (72). A recent report typifies the strategies of testing therapeutic HPV vaccines. Mice were subcutaneously injected with HPV16 E6/E7 expressing TC-1 tumor cells, which expanded to a 9 to 12 mm^2^ area within 10 to 12 days, respectively, and then given a single intramuscular dose of mRNA-lipid nanoparticle vaccines encoding a fusion protein of the type 1 herpes simplex virus glycoprotein D and HPV16 E7 protein (73). The treatments led to regression in 4/5 (80%) and 4/7 (60%) of the respective cohorts; however, there was no indication that the sites of tumor growth were analyzed for residual HPV-positive tumor cells. Thus, our approach to testing established PV lesions in this study more closely represents clinically relevant HPV disease, making our tumor regression results particularly remarkable.

A major strength of this study is the use of pathologist-guided, quantitative tumor tissue analyses of tumor thickness and a battery of biomarkers relevant to PV-induced tumorigenesis. Coupled to robust statistical analyses, our objective quantification of biomarkers reveals clearly that MEK inhibitors accessed the MmuPV1-infected epithelium to suppress p-ERK1/2 levels, basal cell expansion, and tumor thickness/volume without affecting overall epithelial differentiation. Whereas it is most likely that MEK inhibition suppressed tumor growth *via* both blunting ERK-mediated cellular proliferative signals and reducing the transcription of MmuPV1 early genes that promote cell proliferation, the precise contribution of each of these interdependent activities to the overall decreased tumor growth will be challenging to dissect.

The MmuPV1 tumor growth inhibitory effect of the MEK1/2 inhibitor, trametinib, was superior to that of cobimetinib, which preferentially targets the MEK1 isoform. Despite the functional redundancy of MEK1 and MEK2 in many assays (39), our understanding of their distinct activities and differential regulation is expanding. For example, MEK1 contains a site (Thr292) for ERK-mediated feedback phosphorylation whereas MEK2 lacks this site (40, 41). Additionally, several scaffold proteins specifically bind to and enhance the activation of MEK1, but not MEK2 (42–45). MEK1 knockout in mice is embryonic lethal (39, 47), whereas MEK2 knockout animals are viable, fertile, and phenotypically normal (48). MEK2, but not MEK1, phosphorylates the tumor suppressor GCIP (Grap2- and cyclin D1-interacting protein), promotes its degradation, and decreases its tumor suppressor function (46). Our work strongly suggests that inhibition of both MEK1 and MEK2 is crucial for controlling PV disease. However, MEK2 appears to have distinct functions important for MmuPV1 tumor growth that will require futher dissection. There could certainly be other biologicial explanations for the greater potentcy of trametinib.

The ability of MEK1/2 inhibition by trametinib to promote tumor regression became evident only four days post treatment, when the mean tumor volumes in the drug-treated groups were smaller than the group’s mean starting tumor volumes (Fig. 1C, *SI Table 1*). While the tumors in the trametinib-treated groups continued to decrease in size over the next two weeks, the growth reduction seemed to plateau thereafter. This raises important questions as to whether continued trametinib treatment would eventually cure the infections, or whether the drug had reached its maximum effects. It seems likely that ceasing treatment with the drugs prior to complete MmuPV1 clearance would result in regrowth of the tumors, but this has yet to be tested and it would also be informative to do so in the context of functional T cells.

The mechanisms by which the MEK inhibitors exerted their effects in this MmuPV1 infection and tumorigenesis model are consistent, in part, with those observed in other non-PV-related cancer models. In a variety of cancer cell types *in vitro*, trametinib and cobimetinib have been shown to reduce p-ERK1/2, modulate cyclin D1 and p27 levels in a manner aligned with a G1/S cell cycle arrest, and induce caspase-dependent apoptosis (61, 74, 75). Unfortunately, a limitation of our study was the inability to detect nuclear antigens in the lesions due to Versenate treatment for tail decalcification before fixation. Because Sundberg et al. showed similar staining patterns between Ki67 and K5 in MmuPV1-induced tail tumors (56), and we found a correlation between p-ERK1/2 levels and strong K5 staining (r=0.5947, p=0.025; Fig. S5B), we infer that MEK inhibitors suppressed cellular proliferation. Additionally, the strong correlations between p-ERK1/2 levels and MmuPV1 E6/E7 mRNA (r=0.8303, p= 6.85e-5) and between strong K5 staining and MmuPV1 E6/E7 mRNA (r=0.8202, p=9.995e-5) highlight the interdependency of the biomarkers p-ERK1/2, K5 and MmuPV1 E6/E7 mRNA transcription (Fig. S5B, E, I). Future studies evaluating cellular markers of proliferation, apoptosis, and senescence will be needed to clearly delineate the antiproliferative effect of MEK inhibition in mammalian PV tumors.

In our previous study and in this work, the antiviral and tumor suppressive effects of MEK inhibition occurred in the absence of T cells (32). This has important clinical implications for treating PV diseases in those who are immunocompromised (76). For example, people living with AIDS have increased HPV acquisition, prevalence, persistence, high tumor burdens, and cancer progression (77) and would greatly benefit from effective HPV antiviral treatments. Herein, neither T cells nor neutrophils were implicated tumor regression despite reports that immune mediated cell death is an important characteristic of tumor reduction (reviewed in (78)). Whether immune cell populations are involved in the MEK-inhibitor induced regression of PV neoplasia will require further investigation. Importantly, use of MEK1/2 inhibitors as a therapeutic for PV diseases must be tested in the context of functional T cells as MEK signaling is induced by T cell receptor signaling (79). While this could be a disadvantage to tumor clearance, recent evidence indicates that MEK inhibitors can lead to a shift from a Th2 to a Th1 response (80), a reduction of regulatory T cells (81), and reprogramming of CD8+ T cells into memory stem cells with antitumor activities (82), each of which might further benefit treatment of PV diseases. Lastly, we provide strong evidence that MEK inhibitors reduced productive MmuPV1 infection measured by both the reduced levels of the viral capsid protein, L1 in tumors, and the lack of secondary muzzle infections observed in the MEK inhibitor-treated mice. In total, our work strongly suggests that MEK1/2 inhibitors warrant further investigation for treatment of mammalian PV infections *in vivo*; such an approach might also reduce PV transmission to new susceptible epithelial sites or between individuals.

## Materials and Methods

### MmuPV1 infections, tumor growth and treatments

All animal work was approved by the University of New Mexico’s IACUC (protocol #20-201020-HSC). Female Hsd:Athymic Nude-*Foxn1^nu^* outbred mice (Envigo) 6-8 weeks of age were inoculated with MmuPV1 virions (a kind gift from Prof. Paul Lambert, Univ. of Wisconsin-Madison). Specifically, the mice were anesthetized and maintained in anesthesia using 3% isoflurane in O_2_. Prior to inoculation, the tail of each mouse was scarified using a 27.5-gauge needle. To standardize infection area and papilloma growth the scarification was initiated 0.5-cm distal to the tail base and continuing for 1-cm toward the tip. Each mouse tail was superficially scarified by scratching over the epithelial surface 20-30 times as described (83). Scarification and viral inoculations were carried out in a biosafety cabinet. MmuPV1 virions (2×10^8^ viral genome equivalents [VGE] in 2 µl) were applied across the entirety of each wound site using a siliconized pipette tip. The MmuPV1 inocula were allowed to penetrate the wound before returning mice to microisolator (biocontainment) cages. Mice were monitored weekly for weight and the appearance of warts. Tumor volumes were determined with digital calipers (Bel-Art Products) using the formula (length*width*height): length was measured from the sagittal plane, width from the transverse plane, and height was measured from the base to the highest peak. When mice had developed papillomas with a volume of ≥17 mm^3^, the animals were stratified by papilloma size and assigned into five groups with similar average tumor volumes (≈40 mm^3^). Trametinib (Selleckchem) and Cobimetinib (Selleckchem) in vehicle mixture (30% PEG-300 + 5% DMSO + 5% Tween-80 in dH_2_O) were administered by oral gavage and tumor dimensions and body weights were recorded every 48 h. Efficacy was measured as percent tumor growth inhibition (TGI) relative to the vehicle-treated group. TGI was calculated by the equation [1-(*T/C*) x 100], where *T* and *C* represent the mean tumor volumes on day 30 in the MEK inhibitor-treated (T) and vehicle control (C) groups, respectively. A TGI of >50% is considered to be efficacious. On day 30, mice were euthanized by CO_2_ exposure and whole tail sections receiving the MmuPV1 inoculation were excised to include ≈0.5 cm of uninfected length on each side of the wart growth. Tail sections were fixed in 4% paraformaldehyde (PFA) in PBS overnight at 4°C. Thereafter, each tail section was rinsed twice with sterile PBS and incubated with Versenate decalcifying reagent (StatLab) at 4°C until the tail bone became pliable. Tail sections were rinsed with sterile PBS and stored in 70% ethanol at 4°C until paraffin-embedding. Muzzle papillomas were excised and stored in 4% PFA in PBS overnight before washing twice with PBS and storing in 70% ethanol until paraffin-embedding.

### MmuPV1 infections *in vitro*

HaCaT cells were seeded at 10^4^ cells per plate in DMEM with 10% fetal calf serum and allowed to adhere overnight at 37°C in 5% CO_2_. Cells were pretreated for 1h with medium containing vehicle (0.1% DMSO) or 1.0 to 100 nM trametinib prior to and during MmuPV1 exposure at 1000 VGE per cell. Infections proceeded for 48h before harvesting for total DNase I-treated RNA. Following reverse transcription (RT), qPCR was performed using primers to amplify spliced MmuPV1 E1^E4 cDNA and normalized to ribosomal protein 18s mRNA levels as previously described (32, 83, 84).

### Tissue Staining and Immunohistochemistry (IHC)

Formalin-fixed paraffin embedded (FFPE) tail sections and muzzle specimens 4-5 µm in thickness were baked at 60°C for 1 h. The Ventana Discovery platform was used for deparaffinization, rehydration and H & E staining or blocking of endogenous peroxidases for IHC. Antigens were unmasked by boiling in RiboCC buffer (Ventana 760-107; for keratin 10 and myeloperoxidase) for 32 min at 100°C. Antibodies to cytokeratin 10 (abcam ab76318; 1:250), cytokeratin 5 (abcam ab52635; 1:400), and myeloperoxidase (abcam ab208670; 1:1000) were individually diluted in Discovery PSS Diluent (Ventana 760-212), hand applied, and incubated for 16-32 min at 36°C. Antibody incubation was followed by anti-Rb HQ (Ventana 760-4815), anti-HQ HRP (Ventana 760-4820), and DAB CM (Ventana 760-4304) detection. Immunohistochemical detection of p-ERK1/2 (Cell Signaling #4370; 1:500) required the tyramide signal amplification (TSA) biotin system following the manufacturer’s protocol (AKOYA Biosciences #NEL 700A001KT). Slides were counterstained with hematoxylin (Ventana 760-2021) and Bluing solution (Ventana 760-2037) or with Gill No.1 Hematoxylin solution (Sigma-Aldrich #GHS132-1L) followed by acid alcohol and lithium carbonate rinse. Following dehydration, coverslips were mounted using Permount (Fisher Scientific SP15-500). Stained slides were scanned at 20X using the Leica Versa 200 digital slide scanner (Leica Biosystems, Buffalo Grove, IL).

### DNA and RNA *in situ* hybridization (ISH)

FFPE tail sections were stained according to the RNAscope manufacturer’s protocol (2.5 HD Detection Kit-Red, Advanced Cell Diagnostics [ACD]) using probes to detect MmuPV1 E6/E7 RNA (ACD). DNA ISH employed probes to the noncoding MmuPV1 LCR Probe (ACD) using the chromogenic DNAscope HD Kit. Slides were counterstained with 50% Gill’s hematoxylin solution number 1 (G Biosciences), mounted using EcoMount (BioCare Medical), and digitally imaged as above.

### Immunofluorescence

FFPE tail sections were deparaffinized, rehydrated, and subjected to heat-induced antigen retrieval for 30 min using pH 9 target retrieval solution (Dako). Endogenous peroxidases were blocked with 3% hydrogen peroxide in methanol for 10 min. Tissues were blocked with 5% goat serum in PBS containing 0.1% Tween-20 (PBS-T) for 1 h at room temperature, then incubated at 4°C overnight with rabbit sera against MmuPV1 L1 (a kind gift form Chris Buck, NIH; 1:2500) and chicken anti-keratin 14 (BioLegend 905301; 1:500) diluted in 5% goat serum in PBS-T. The sections were then washed three times in PBS-T followed by incubation with secondary antibodies (1:500) Alexa fluor (AF)-488 goat anti-rabbit IgG (ThermoFisher A32731) and AF-647 goat anti-chicken IgG (ThermoFisher A32933) for 1 h at room temperature. Coverslips were mounted using ProLong Diamond Antifade DAPI Mountant (Thermo Fisher). Images were acquired using a Leica TCS SP8 confocal microscope with fixed data acquisition settings. L1-positive foci were counted using Huygens essential software object analysis tool (Scientific Volume Imaging, The Netherlands, http://svi.nl).

### Morphometric Tissue Analysis

Tissue morphometry was performed on digitally acquired images from all tumors using HALO software (Indica, Albuquerque, NM), under the supervision of a board-certified veterinary pathologist (Kusewitt). The thickness of the epithelium within each lesion was determined using H&E sections. The linear length of the lesion was measured, and HALO was trained to recognize all epithelial elements lying above the dermis, then the area of epithelium within the lesion was determined. The epithelium within each region was selected using the bucket tool; the selection was refined using the line and scissors tools to include all nucleated cells and to eliminate artifacts. Within the identified epithelium, stroma was identified and eliminated from consideration. Epithelial thickness was expressed as the ratio of epithelial area to lesion length (mm). Using the CytoNuclear algorithm, individual epithelial cells were identified and cytoplasmic staining for each marker was categorized as negative, low, medium, or high. The total number of cells examined was enumerated and the percentage of epithelial cells staining at each intensity was determined (e.g., *SI Appendix* Fig. S3A-C). As the ISH staining lacked sufficient nuclear demarcation, total areas positive for ISH signals were determined. Neutrophil infiltration was quantified by selecting all lesional epithelium and underlying dermis by hand. The percentage of that area occupied by myeloperoxidase-positive neutrophils was determined using HALO software to identify and quantify the area of positive staining. IHC stained slides were excluded from analysis (no more than one per treatment group) if they lacked clear evidence of tail features that ensured the tumor was adequately represented.

### Statistical Analyses

Morphometry analyses included data from 5-6 tumors per group subjected to one-way analysis of variance (ANOVA) with Tukey’s multiple comparison test. RT-qPCR data were analyzed by one-way ANOVA with Dunnett’s multiple comparisons test. Biomarker level correlation coefficients (*r*) were calculated from morphometry data using Pearson’s correlation analysis and computing a two-tailed *p* value. Tumor measurements among the five treatment groups were analyzed at each time point using the generalized least square (GLS) method to account for the unequal variances among the groups (*SI Appendix* Table 1). Differences among tumor groups at 30 days post treatment were evaluated by ANOVA with the Tukey HSD (honest significant difference) pairwise comparisons (*SI Appendix* Table 2). The rates of tumor volume changes were estimated and compared using the linear mixed-effects model (LMM) analysis (*SI Appendix* Tables 3 and 4).

## Acknowledgments

This study was supported by the American Cancer Society Mission Boost Grant MBG-18-207-01-COUN (M.A.O.); and the US National Institutes of Health Grants: R21DE028652 (M.A.O.); T32AI07538, F31CA232536, UNMCCC Matching Support (A.J.L.); P30CA118100 (M.A.O.). We acknowledge support by the following University of New Mexico Comprehensive Cancer Center shared resources (NCI 2P30CA118100): Human Tissue Repository and Tissue Analysis, Animal Models, Biostatistics.

We are grateful to Dr. Paul Lambert and members of his lab for generously providing infectious stocks of MmuPV1, to Dr. Nagayasu Egawa for sharing his protocols for mouse tail scarification and infection, and Dr. Christopher Buck for sharing rabbit sera against MmuPV1 L1.

## Abbreviations

AP-1: Activator protein-1
AF: Alexa fluor
ANOVA: analysis of variance
DMSO: dimethyl sulfoxide
FFPE: formalin-fixed paraffin embedded
GLS: generalized least square
HPVs: human papillomaviruses
HPV: human papillomavirus
IHC: immunohistochemistry
IC_50_: inhibitory concentration 50
ISH: *in situ* hybridization
K5: keratin 5
K10: keratin 10
LMM: inear mixed-effects model
LCR: long control region
DMEM: Dulbecco’s modified Eagle’s medium
MmuPV1: murine papillomavirus type 1
PFA: paraformaldehyde
PBS: phosphate-buffered saline
TGI: tumor growth inhibition
VGE: viral genome equivalents

## Supplementary Information for

**This PDF file includes:**

Figures S1 to S5

Tables S1 to S4

**Other supplementary materials for this manuscript include the following:**

None

**Fig. S1.**
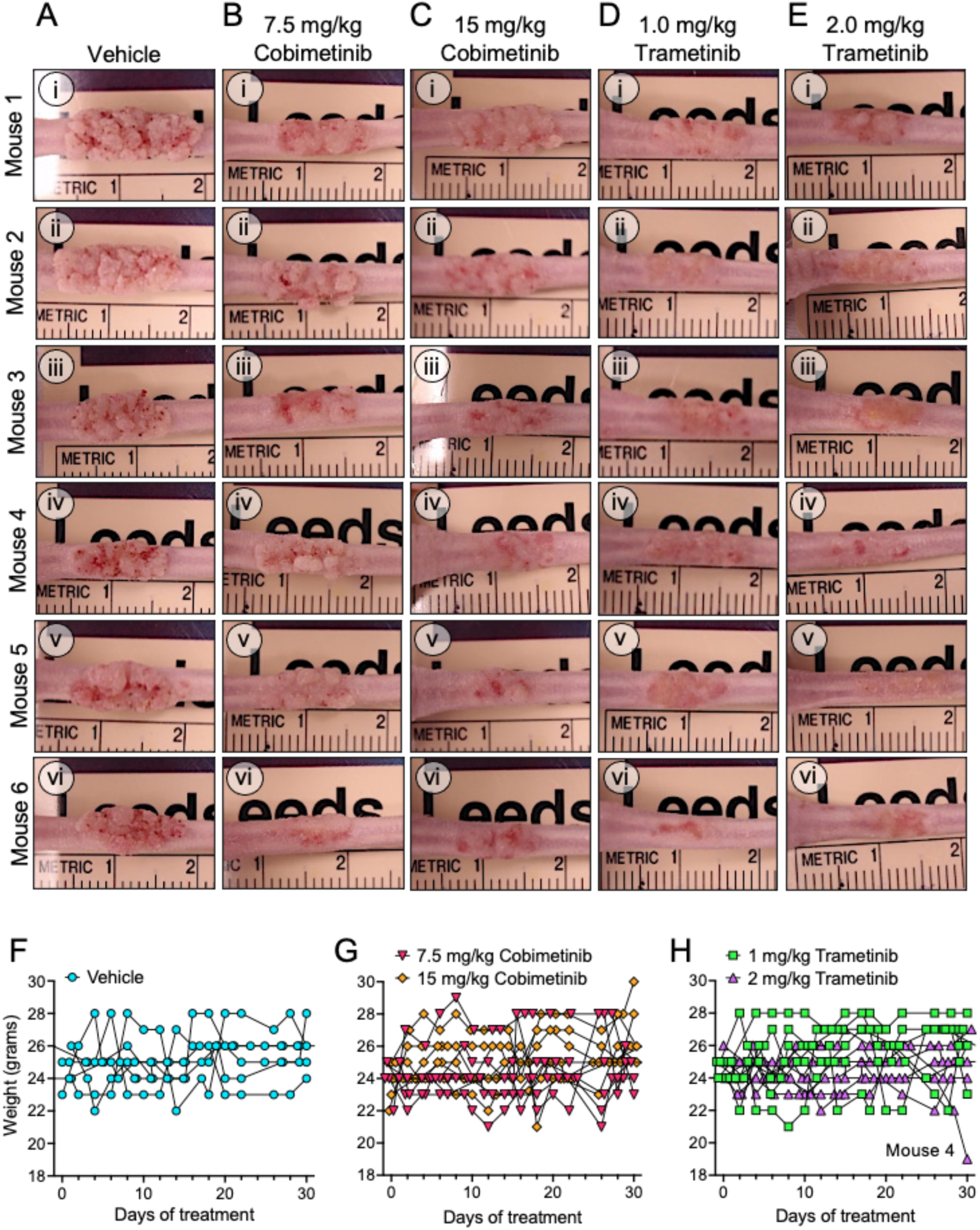
Visual phenotypes of MmuPV1-induced papillomas at day 32 post treatment (prior to harvest). MmuPV1-induced papillomas after treatment with vehicle (A), 7.5 mg/kg cobimetinib (B), 15 mg/kg cobimetinib (C), 1 mg/kg trametinib (D), or 2 mg/kg trametinib (E). The labels at the left identify animal numbers in each treatment group. (F-H) Mouse weights during treatment period. Note that only one animal (mouse 4 of the 2 mg/kg trametinib treatment arm) showed substantial weight loss (>20% of starting weight).

**Fig. S2.**
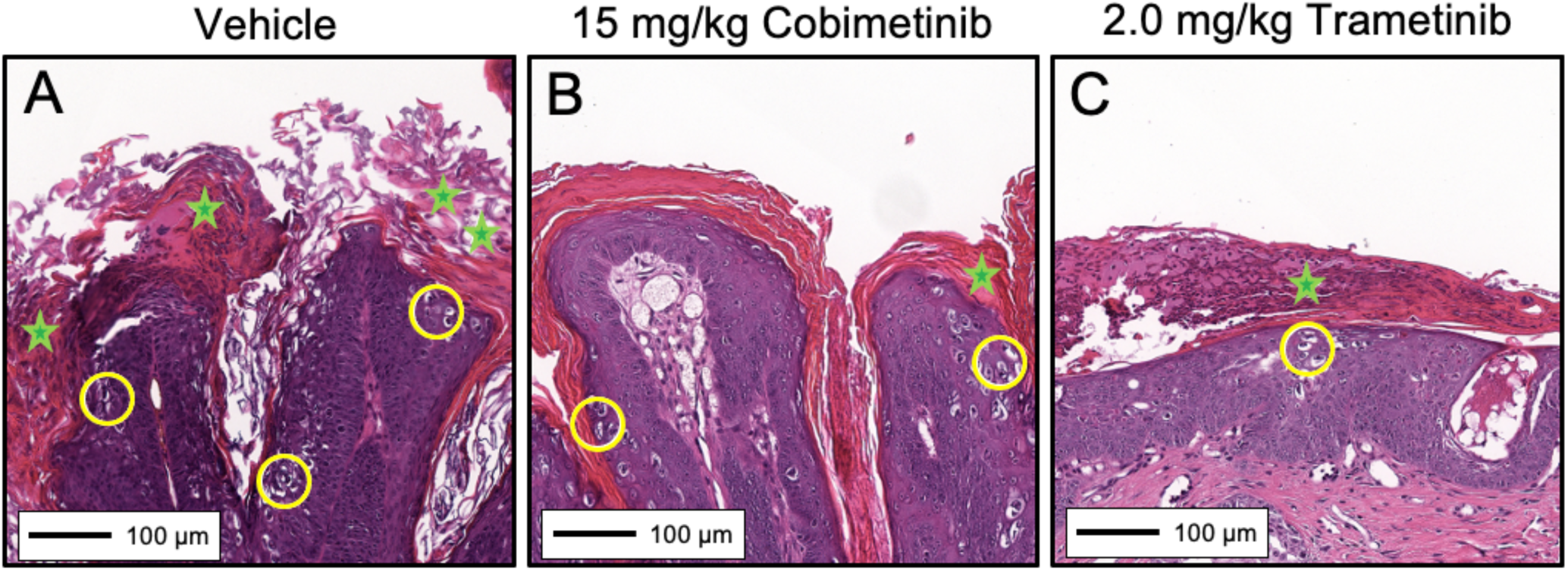
Histologic features of MmuPV1-induced papillomas. High-magnification images of H&E-stained papillomatous tissue sections from animals treated with vehicle (A), 15 mg/kg cobimetinib (B) or 2.0 mg/kg trametinib (C). Regions of hyperkeratosis are denoted with green stars. Koilocytes, a hallmark of papillomavirus infections, are encircled in yellow.

**Fig. S3.**
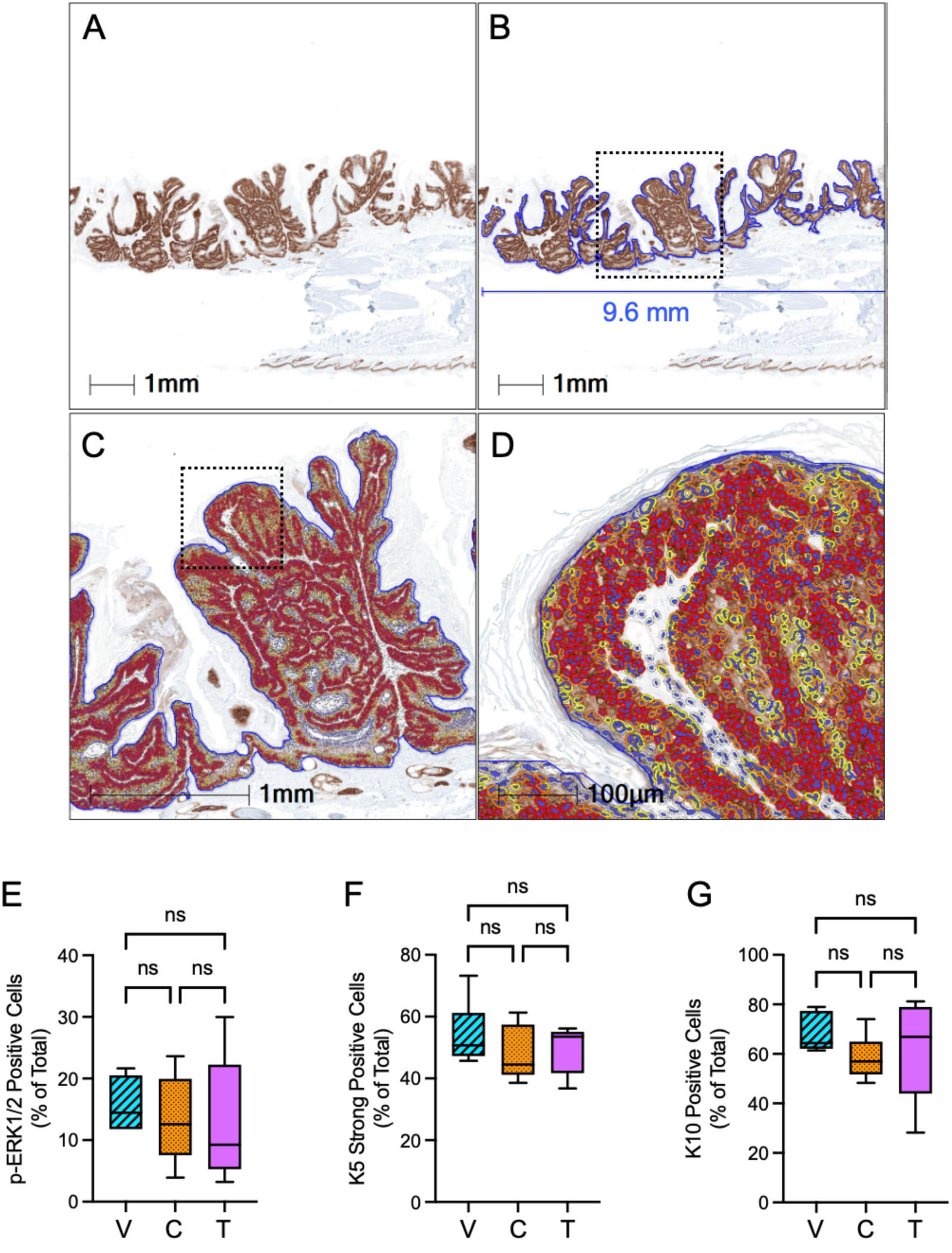
Example of pathologist-guided morphometric analysis using a HALO algorithm to identify and quantify the areas of strong positive K5 staining and comparisons of % biomarker signals amongst the tumor groups. (A-B) Tumor-bearing areas were selected (blue) and defined as nuclei-containing epithelium; the boxed area in B was subject to increased magnification in panel C. (C) Using the CytoNuclear algorithm, individual epithelial cells were identified *via* marking nuclei blue and cytoplasmic staining for each marker was categorized as negative (clear), low (yellow), medium (orange), or high (red). (D) Increased magnification of the boxed area in panel C. Because cells gradually loose K5 expression as they begin to differentiate and express K10, only the strong K5 positive cells were enumerated. (E-G) Summary of morphometric analyses for the percentage of MmuPV1-induced tumors cells expressing the cellular proteins p-ERK1/2 (E), strong K5 (F), and K10 (G) for mice treated with vehicle (V), 15 mg/ml cobimetinib (C), 2 mg/kg trametinib (T). Data were analyzed using one-way ANOVA with Tukey’s multiple comparison test (ns = not significant).

**Fig. S4.**
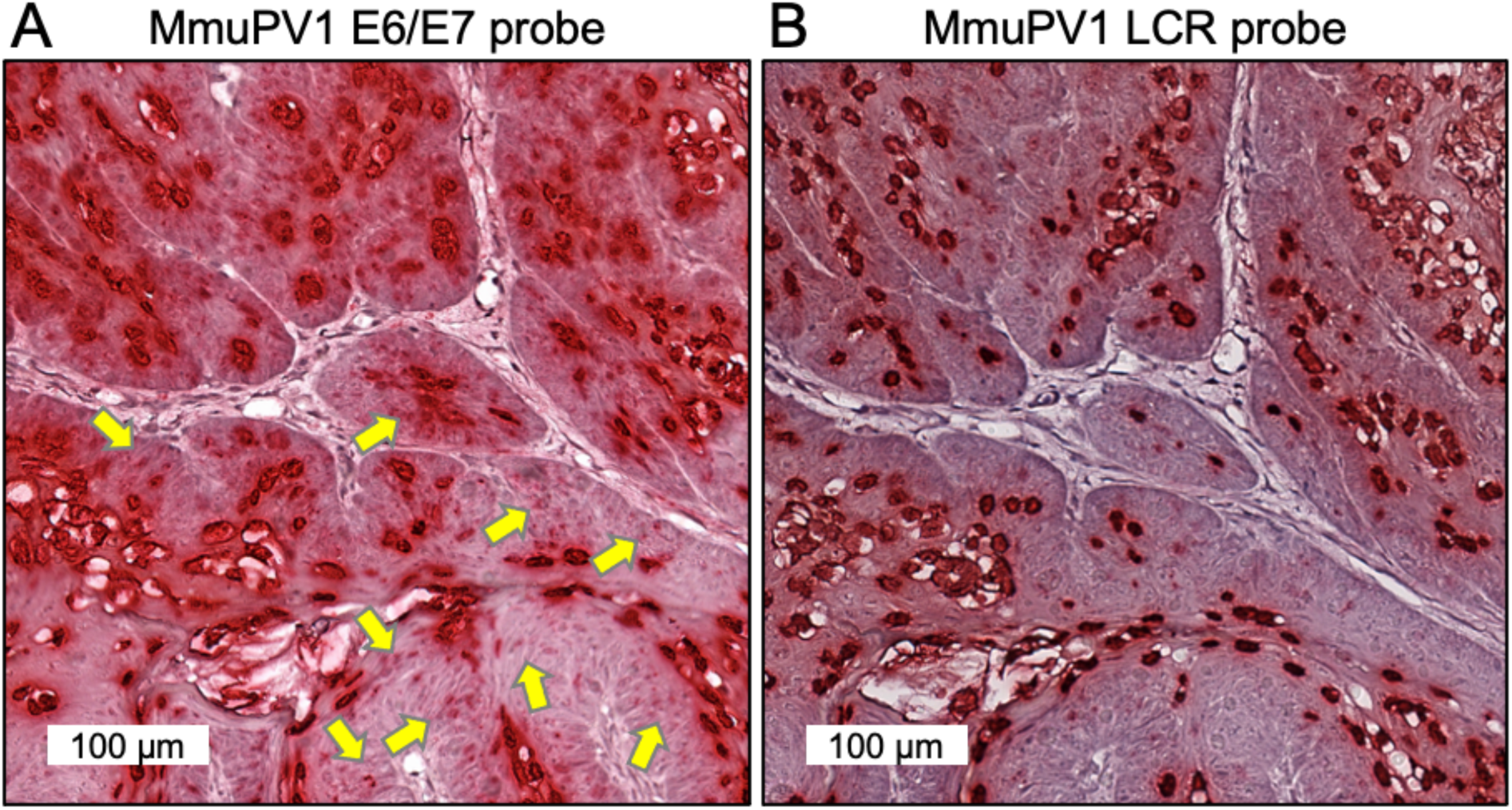
Magnified images of ISH using probes specific for the MmuPV1 E6/E7 coding region (A) and the noncoding genomic LCR segment (B) in adjacent tumor sections from a vehicle-treated mouse. Note that both probes strongly stained nuclei in koilocytes and suprabasal cells likely to be amplifying viral genomes. Whereas the E6/E7 probes demonstrated cytoplasmic punctate signals expected for viral mRNAs (A, yellow arrows), the LCR probe produced little cytoplasmic signal (B).

**Fig. S5.**
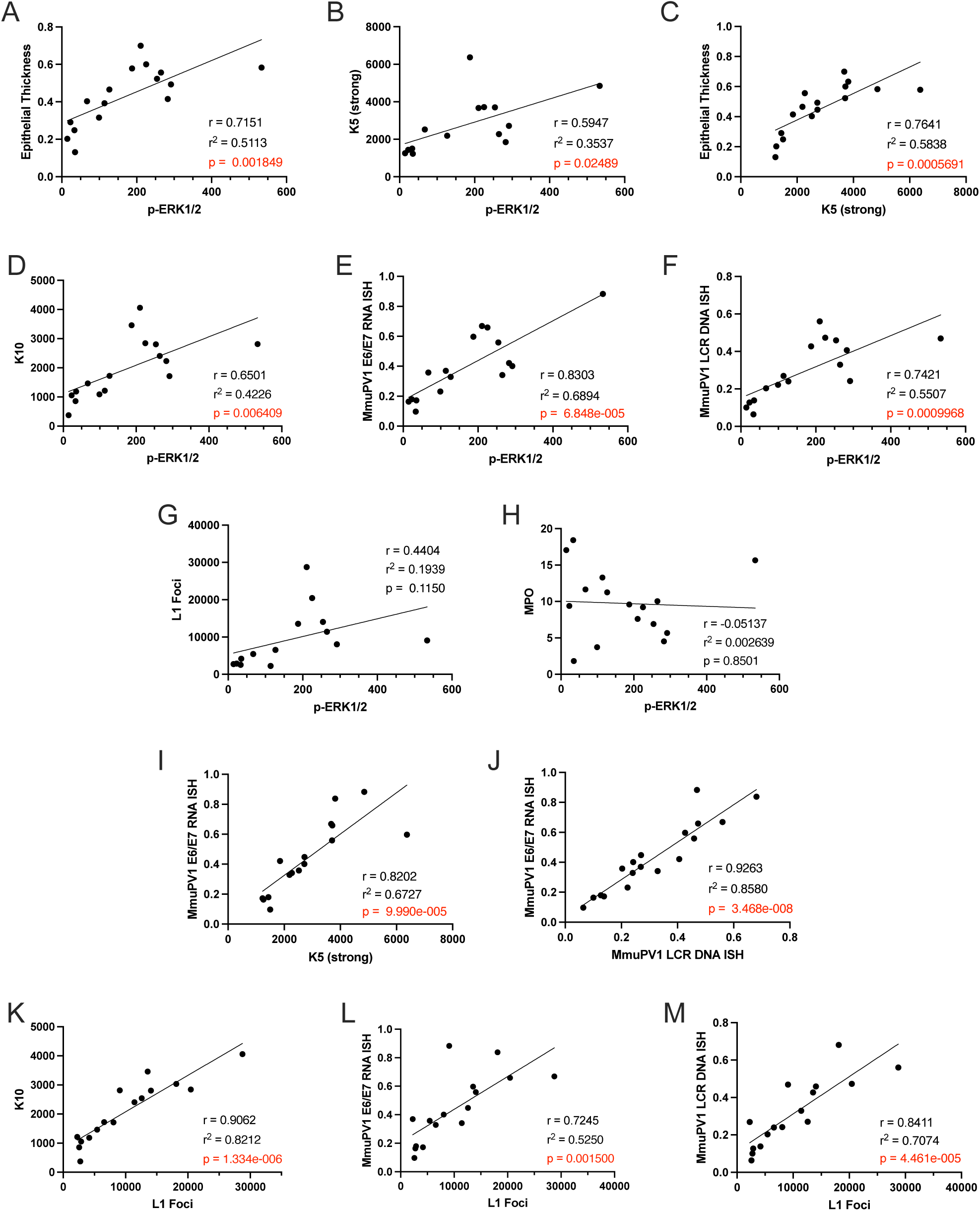
MEK inhibitor reduced tissue levels of p-ERK1/2 correlate with cellular and viral biomarkers in MmuPV1-induced tail tumors. Scatter dot plot correlations analyses between selected biomarkers were calculated using Pearson correlation coefficient (r) analysis and a two-tailed test (p values). P values that reached statistical significance are shown in red.

**SI Table 1.** Mean and Standard Error (SE) of Tumor Volumes (mm^3^)

| **Time**<br>**(days)** | **Vehicle**<br>**Mean (SE)** | **7.5 mg/kg**<br>**Cobimetinib**<br>**Mean (SE)** | **15 mg/kg**<br>**Cobimetinib**<br>**Mean (SE)** | **1.0 mg/kg**<br>**Trametinib**<br>**Mean (SE)** | **2.0 mg/kg**<br>**Trametinib**<br>**Mean (SE)** | **p-value<sup>a</sup>** |
| --- | --- | --- | --- | --- | --- | --- |
| 0 | 39.2 (8.1) | 39.8 (6.3) | 40.8 (6.3) | 39.3 (8.0) | 41.6 (7.5) | 0.998996 |
| 2 | 46.8 (7.1) | 42.1 (5.4) | 48 (11.6) | 40.1 (6.1) | 46.1 (7.5) | 0.906252 |
| 4 | 56.1 (9.0) | 42.6 (5.8) | 41.9 (7.9) | 31.1 (5.5) | 37.8 (8.7) | 0.20922 |
| 6 | 61.2 (12.8) | 47.0 (4.4) | 41.1 (6.0) | 40.9 (8.3) | 34.3 (6.3) | 0.27487 |
| 8 | 65.8 (11.6) | 41.5 (7.5) | 33.5 (9.2) | 30.9 (5.9) | 25.5 (4.3) | 0.041988 |
| 10 | 65.8 (11.2) | 34.9 (5.9) | 37.8 (8.3) | 29.5 (5.1) | 26.0 (5.4) | 0.069824 |
| 12 | 78.9 (12.2) | 36.9 (4.6) | 35.5 (9.3) | 29.1 (7.7) | 24.5 (6.0) | 0.023779 |
| 14 | 84.5 (10.6) | 40.3 (5.7) | 36.0 (9.2) | 24.1 (4.0) | 25.9 (6.2) | 0.001759 |
| 16 | 82.9 (16.0) | 38.5 (5.3) | 35.7 (11) | 23.5 (4.4) | 20.9 (4.4) | 0.005515 |
| 18 | 92.3 (16.0) | 34.0 (6.6) | 32.7 (9.5) | 30.3 (3.7) | 21.8 (4.3) | 0.016097 |
| 20 | 80.7 (17.1) | 38.3 (6.9) | 39.0 (10.7) | 24.8 (5.1) | 22.4 (3.7) | 0.015508 |
| 22 | 84.7 (17.3) | 43.2 (6.2) | 40.0 (10.0) | 24.4 (4.5) | 20.7 (6.1) | 0.004064 |
| 24 | 95.2 (22.1) | 44.4 (7.3) | 37.7 (9.8) | 22.5 (2.6) | 21.4 (4.5) | 0.004354 |
| 28 | 109.2 (19.2) | 51.4 (10.7) | 36.4 (8.8) | 22.6 (4.6) | 23.2 (5.8) | 0.002729 |
| 30 | 101.7 (19.1) | 42.4 (8.0) | 45.4 (13.2) | 25.0 (4.4) | 25.2 (4.8) | 0.00718 |

<sup>a</sup> p-value for the significance of the difference in tumor volume among the five groups obtained using GLS to account for unequal variances.

**SI Table 2.**
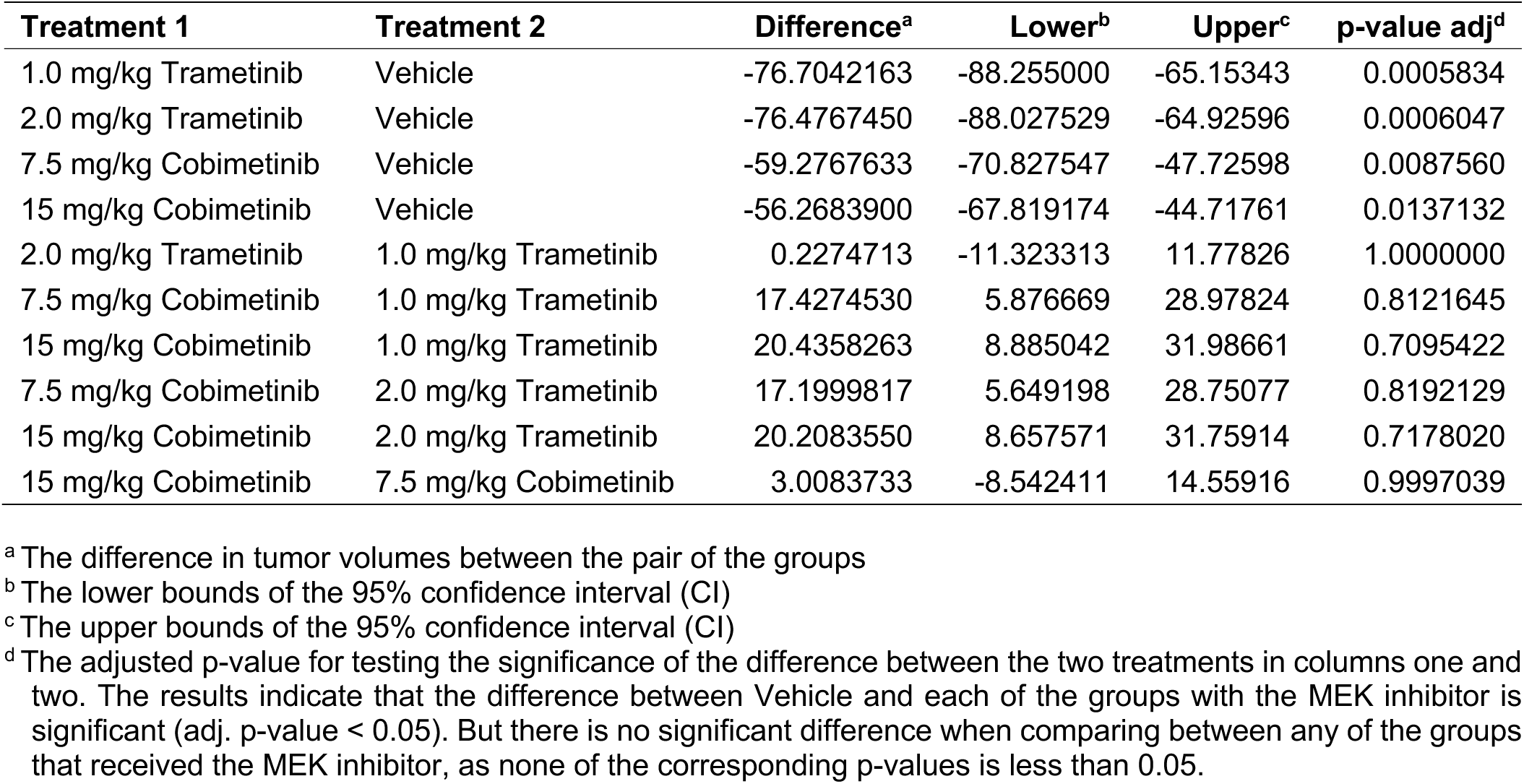
Pairwise Comparisons of Final Tumor Volumes Among Treatment Groups at 30 Days of Treatment.

| **Treatment 1** | **Treatment 2** | **Difference<sup>a</sup>** | **Lower<sup>b</sup>** | **Upper<sup>c</sup>** | **p-value adj<sup>d</sup>** |
| --- | --- | --- | --- | --- | --- |
| 1.0 mg/kg Trametinib | Vehicle | -76.7042163 | -88.255000 | -65.15343 | 0.0005834 |
| 2.0 mg/kg Trametinib | Vehicle | -76.4767450 | -88.027529 | -64.92596 | 0.0006047 |
| 7.5 mg/kg Cobimetinib | Vehicle | -59.2767633 | -70.827547 | -47.72598 | 0.0087560 |
| 15 mg/kg Cobimetinib | Vehicle | -56.2683900 | -67.819174 | -44.71761 | 0.0137132 |
| 2.0 mg/kg Trametinib | 1.0 mg/kg Trametinib | 0.2274713 | -11.323313 | 11.77826 | 1.0000000 |
| 7.5 mg/kg Cobimetinib | 1.0 mg/kg Trametinib | 17.4274530 | 5.876669 | 28.97824 | 0.8121645 |
| 15 mg/kg Cobimetinib | 1.0 mg/kg Trametinib | 20.4358263 | 8.885042 | 31.98661 | 0.7095422 |
| 7.5 mg/kg Cobimetinib | 2.0 mg/kg Trametinib | 17.1999817 | 5.649198 | 28.75077 | 0.8192129 |
| 15 mg/kg Cobimetinib | 2.0 mg/kg Trametinib | 20.2083550 | 8.657571 | 31.75914 | 0.7178020 |
| 15 mg/kg Cobimetinib | 7.5 mg/kg Cobimetinib | 3.0083733 | -8.542411 | 14.55916 | 0.9997039 |

<sup>a</sup> The difference in tumor volumes between the pair of the groups
<sup>b</sup> The lower bounds of the 95% confidence interval (CI)
<sup>c</sup> The upper bounds of the 95% confidence interval (CI)
<sup>d</sup> The adjusted p-value for testing the significance of the difference between the two treatments in columns one and two. The results indicate that the difference between Vehicle and each of the groups with the MEK inhibitor is significant (adj. p-value < 0.05). But there is no significant difference when comparing between any of the groups that received the MEK inhibitor, as none of the corresponding p-values is less than 0.05.

**SI Table 3.** Estimates for Rates of Change of Tumor Volumes (Slope) SI Table 3 shows the estimates for the rate of change (mm^3^/day), the slopes of the linear equations of the five groups, along with the SE, DF, t-value and p-value. The t-test here investigates whether the corresponding slopes are zero. The p-values in the first three rows are all smaller than 0.05, suggesting that the rate of change of tumors in the vehicle-treated mice and of tumors in mice treated with trametinib significantly differ from zero. The tumor volumes in vehicle-treated mice significantly increased over time (because the corresponding slope is positive), and that of groups treated with trametinib significantly decreased over time (as the corresponding slopes are negative). The tumor volumes in groups treated with cobimetinib have the smallest estimates for the slopes (in the sense of the absolute value), and the corresponding p-values are very large, indicating that the rate of change is not significantly different from zero (i.e., the tumor volumes, on average, did not have a significant change over time).

| Group | Estimate<sup>a</sup> | SE<sup>b</sup> | DF<sup>c</sup> | t-value<sup>d</sup> | p-value<sup>e</sup> |
| --- | --- | --- | --- | --- | --- |
| Vehicle | 1.849 | 0.615 | 445 | 3.01 | 0.0028 |
| 1 mg/kg Trametinib | -0.4739 | 0.2262 | 445 | -2.09 | 0.0368 |
| 2 mg/kg Trametinib | -0.5325 | 0.254 | 445 | -2.1 | 0.0366 |
| 7.5 mg/kg Cobimetinib | 0.1011 | 0.2795 | 445 | 0.36 | 0.7178 |
| 15 mg/kg Cobimetinib | 0.1168 | 0.3679 | 445 | 0.32 | 0.7509 |

<sup>a</sup> Estimates for the rate of change (mm<sup>3</sup>/day) derived from the slopes of the linear equations
<sup>b</sup> The standard error (SE)
<sup>c</sup> The degree of freedom (DF)
<sup>d</sup> The value of t-test (t-value) where the t-test investigates whether the corresponding slopes are zero.
<sup>e</sup> The p-value investigates whether the rate of change significantly differs from zero.

**SI Table 4.** Difference in Rates of Change of Tumor Volumes (Slope) SI Table 4 shows the analysis of the individual tumor rates of change for each treatment group using the linear mixed-effects model (LMM). Based on the fitted LMM, we estimated the differences in the rate of change for comparisons of interest. The first four rows provide the estimates for the difference in the rate of change between the Vehicle and each MEK inhibitor treatment group. The fact that the p-values are smaller than 0.05 indicates that the rate of change differs significantly between each pair of the groups. The fifth row shows the comparison between the Vehicle and overall treatment group (the union of all the treatment groups), and a small p-value implies that the difference in the rate of change is significant. These results provide statistical evidence for the considerable effect of MEK inhibitors on tumor growth. In rows six and seven, we compared the dose effect on the rate of change among the same drug, and the large p-values indicate that the dose effect is insignificant. Lastly, we compared the difference in the rate of change between the two drugs regardless of the dose. The result indicates a significant difference in the rate of change between those treated with the two drugs (p-value = 0.0334).

| Difference in Rate of Change | Estimate<sup>a</sup> | SE<sup>b</sup> | DF<sup>c</sup> | t-value<sup>d</sup> | p-value<sup>e</sup> |
| --- | --- | --- | --- | --- | --- |
| Between Vehicle and 1 mg/kg Trametinib | 2.3229 | 0.6553 | 445 | 3.54 | 0.0004 |
| Between Vehicle and 2 mg/kg Trametinib | 2.3816 | 0.6654 | 445 | 3.58 | 0.0004 |
| Between Vehicle and 7.5 mg/kg Cobimetinib | 1.7479 | 0.6756 | 445 | 2.59 | 0.01 |
| Between Vehicle and 15 mg/kg Cobimetinib | 1.7322 | 0.7167 | 445 | 2.42 | 0.016 |
| Between Vehicle and Overall Treatment | 2.0462 | 0.6315 | 445 | 3.24 | 0.0013 |
| Between Two Doses of Trametinib | 0.05866 | 0.3402 | 445 | 0.17 | 0.8631 |
| Between Two Doses of Cobimetinib | -0.01576 | 0.462 | 445 | -0.03 | 0.9728 |
| Between Trametinib and Cobimetinib | -0.6122 | 0.2869 | 445 | -2.13 | 0.0334 |

<sup>a</sup> Estimates for difference in the rate of change (mm<sup>3</sup>/day) between the two groups of interest obtained based on the LMM
<sup>b</sup> The standard error (SE)
<sup>c</sup> The degree of freedom (DF)
<sup>d</sup> The value of t-test (t-value) where the t-test investigates whether the corresponding differences are zero.
<sup>e</sup> The p-value investigates whether the rate of change significantly differs between the groups of interest.

